# Sand flies around Tona, an artificial water reservoir in Bucaramanga (Santander, Colombia): diversity, spatiotemporal frequency, and efficacy of four sampling methods

**DOI:** 10.1101/2024.01.18.576224

**Authors:** Laura Rengifo-Correa, Ruth M Castillo, Juliana Cuadros, Eunice A B Galati, Jonny E Duque

**Affiliations:** Center of Research in Tropical Diseases – CINTROP. Health Faculty, Medicine School, Basic Sciences Department, University Industrial of Santander, Santander, Colombia; Department of Epidemiology. School of Public Health. University of Sao Paulo.

**Keywords:** Phlebotomine, new records, *Pintomyia youngi*, sticky traps, vectors in water reservoirs

## Abstract

Characterizing the sand fly diversity is crucial for an effective design of control strategies for Leishmaniasis. We aimed to characterize the sand fly biodiversity around the Tona Reservoir (Santander, Colombia) and to compare the performance of four sampling methods. A sampling of insects was performed in 2017 in the most preserved and least preserved areas close to the Tona Reservoir, using Torre Vigia-UIS and HomeTrap-UIS, two sticky traps previously designed by our working group, CDC, and Bg-Sentinel traps. We collected 268 Phlebotominae specimens, most with CDC (47.8%) and Torre Vigia-UIS (30.2%) traps. Some specimens (47%) could not be determined because of their preservation status; these samples came mostly from the sticky traps. We found 16 sand fly morphospecies, of which 12 were determined to species level. Here, we report two new records for Santander: *Pintomyia youngi* (Feliciangeli and Murillo, 1987) and *Lutzomyia ceferinoi* (Ortíz and Álvarez, 1963). We also collected some confirmed vectors of *Leishmania*: *Pi. youngi*, *Lutzomyia gomezi* (Nitzulescu, 1931), and *Lu. longipalpis* (Lutz and Neiva, 1912). The highest diversity was collected in the most disturbed area (15 spp.), and in the rainy season (April, 12 spp.). *Pintomyia youngi* distribution was broad through the Tona Reservoir in all the sampled periods, and we suggest tracking it to infer leishmaniasis risk in the Tona Reservoir. Torre Vigia-UIS seems a valuable tool for vector control, but we do not recommend it for biodiversity studies.

## 1. Introduction

Around 12 million people from 98 countries suffer from leishmaniasis (Alvar et al. 2012), a group of vector–borne diseases caused by obligate intracellular parasitic protozoans of the genus *Leishmania* Ross (Kinetoplastea: Trypanosomatida). Leishmaniasis has broad clinical manifestations, but the most common are cutaneous and visceral forms due to their incidence or lethality (Hong et al. 2020). There are an estimated 1.3 million new leishmaniasis cases annually, of which around 0.7 to 1.2 million are cutaneous and 0.2 to 0.4 million are visceral cases (Alvar et al. 2012). Cutaneous leishmaniasis is the most widespread type and, although usually self-healing, causes long–term repercussions such as face scaring and mutilation of up to 40 million people (Bailey et al. 2017). Visceral leishmaniasis is the most severe form of these diseases, affecting human morbidity and killing more people than most parasitic diseases – ≈50,000 people per year–(Oryan and Akbari 2016, Wamai et al. 2020). Preventing leishmaniasis through vector control is desirable because there is no vaccine and available treatments are not very effective and have many undesirable secondary effects (Stockdale and Newton 2013, Wilson et al. 2020).

Around 10% of sand fly species (Diptera: Psychodidae: Phlebotominae) have been implicated as vectors of *Leishmania* spp. (Oryan and Akbari 2016). Each sand fly species has particular habitat preferences and responds differently to landscape changes (Lewis 1974, Ready 2013, Akhoundi et al. 2016). To design effective control strategies for vectors, it is crucial to characterize the sand fly diversity species and their relationship with their habitats.

Some sand flies of medical relevance can spread from their natural tropical forest or semi-arid habitats to anthropic habitats. A change in the sand fly diversity is a common consequence of habitat transformation because these insects are directly influenced by seasonal variations and new environmental conditions such as deforestation (Oryan and Akbari 2016). One worry with artificial water reservoirs and water reservoir construction is the impact generated on landscapes due to modifying original habitats. The expected consequences of such construction are microclimatic variations in area, displacement, and species loss (Winton et al. 2019). Despite the justified need to collect water for basic human needs, constructing artificial water reservoirs and dams also provides conditions for the spread of vector-borne diseases (Bradley 2012).

Exploration of sand fly diversity is necessary to provide a preliminary inference of leishmaniasis’ epidemiological potential in areas close to new water reservoirs. Diversity studies might be enriched by using multiple sampling methods because some might be more efficient than others. Also, comparative studies allow for the evaluation of new sampling methods in the field. It is worth mentioning that sand fly diversity *per se* is not correlated with leishmaniasis risk because disease transmissivity involves not only vector but also host diversity and climatic context (González-Salazar 2012, González-Salazar et al. 2013). In the epidemiological context of leishmaniasis, recording confirmed vectors in each area is more important than diversity *per se*, because these species contribute more to an area’s epidemiology than non-confirmed vectors. Then, diversity studies of sand flies with the perspective of medical relevance must also include a description of the studied species’ composition.

However, providing a rigorous taxonomic treatment of sand flies is a challenging task because of the group’s taxonomic complexity, particularly in megadiverse countries where sand fly diversity paradoxically contrasts with the low number of specialists working in these areas. Phlebotominae has about 1060 species, of which 555 are in the Americas (Galati and Rodrigues 2023). For Colombia –a megadiverse country– 199 species are recorded, and 18 of them are of medical relevance (Ferro et al. 2015, Bejarano and Estrada 2016). This country is among the nine with the highest incidence of cutaneous leishmaniasis worldwide (Alvar et al. 2012), and its vectors of leishmaniasis are widely distributed, with 30 of its 32 departments recorded (Ferro et al. 2015). Santander department has a significantly high amount of sand fly diversity for the country (51 species), with approximately a quarter of Colombiás sand fly species (Bejarano and Estrada 2016, Gómez-Vargas and Zapata-Úsuga 2022). Records of these species in Santander mainly come from wet and dry rainforest ecosystems (Gómez-Vargas and Zapata-Úsuga 2022). Santander has nine species confirmed as leishmaniasis vectors: *Lutzomyia gomezi* (Nitzulescu, 1931), *Lu. hartmanni* (Fairchild and Hertig, 1957), *Lu. lichyi* (Floch and Abonnenc, 1950), *Lu. longipalpis* (Lutz and Neiva, 1912), *Nyssomyia etrapidoi* (Fairchild and Hertig, 1952), *Ny. yuilli* Young and Porter, 1972, *Psychodopygus panamensis* (Shannon, 1926), *Pintomyia evansi* (Núñez-Tovar, 1924), *Pi. ovallesi* (Ortíz, 1952) (Bejarano and Estrada 2016, Gómez-Vargas and Zapata-Úsuga 2022). Here, we aimed to characterize the biodiversity of sand flies in the foothills area of a recently built artificial water reservoir, “Tona”, located in a tropical wet forest in the Santander Department of northern Colombia. Also, we compared the performance of two sticky traps previously designed by our working group (Torre Vigia-UIS and HomeTrap-UIS), with CDC and Bg-Sentinel traps.

## 2. Materials and methods

### 2.1 Ethics statement

We obtained permission for the collection of sand flies from the Autoridad Nacional de Licencias Ambientales of Colombia (Permission framework for the collection of specimens of wild species of biological diversity for non-commercial scientific research purposes, Ref. No. IDB0398-ANLA).

### 2.2 Area of study

The study was conducted in the Retiro Grande village (7°09’ – 8°09’N and 73°05’ – 74°05’W) in the rural area of Bucaramanga, the capital city of Santander. The Santander department is located in the northwestern part of Colombia. The Andean Mountains mark its orography from the western area, followed by a plain formation in the eastern area (Figure 1). The altitude of our area of study oscillates from 800 to 943 m above sea level. This place is a wet forest or bh-PM (Rincón et al. 2019), with a mean annual temperature of 23°C, 1317 mm/yr rainfall, and two rainy seasons: April to June and September to November. In 2016, the Acueducto Metropolitano of Bucaramanga built Tona’s artificial water reservoir in the Retiro Grande village. This reservoir has 17.6 million cubic meters of water, regulating a flow of more than 1000 L/sec to Bucaramanga, Girón, and Floridablanca cities (amb 2017). The Tona Reservoir has preserving (pre-Tona Reservoir) and intervening (intra– and post-Tona Reservoir) areas. The post-reservoir zone is the final water reservoir area; it has a road and a tunnel connecting the Tona municipality with localities near the water reservoir. Human settlements are not allowed inside the vicinity of Tona Reservoir, but the water reservoir staff integrates the place’s float population. In spite of this, some sparse, non-regulated houses still exist close to the Tona Reservoir. The densest urban area neighborhood, Chitota, is approximately five kilometers from the water reservoir.

**Figure 1.**
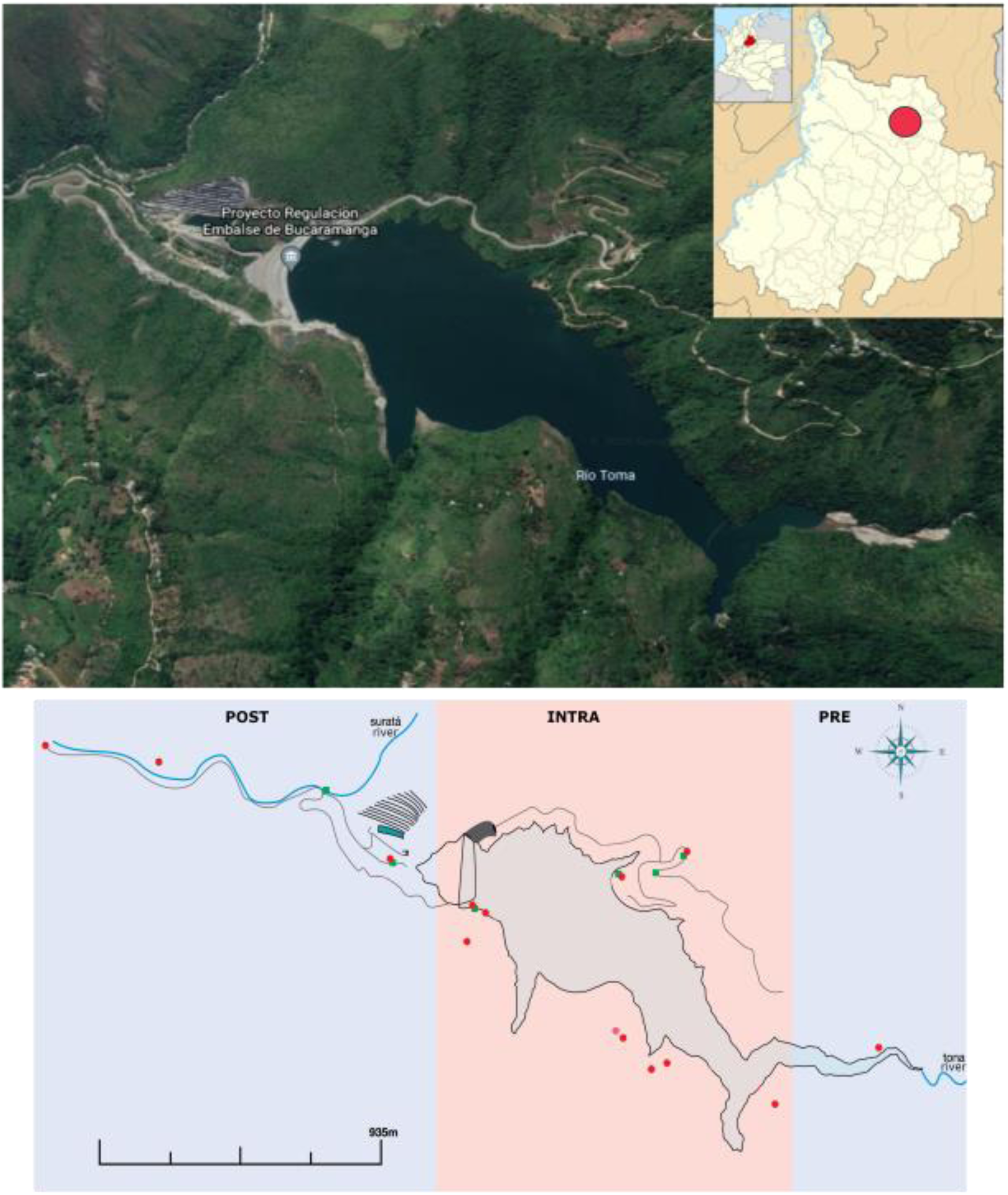
(a) The artificial water reservoir Tona’s geographic location (Santander, Colombia). (b) Sampling points through post-, intra-, and pre-Tona reservoir.

### 2.3 Spatio-temporal sampling

We carried out monthly entomological surveillance from February to October 2017. Sand flies were collected from 14 sampling points, which we selected based on the transition features of the water reservoir (pre-, intra-, and post-reservoir). Due to the absence of published sand fly reports, traps were installed in different places according to the features of the collection points (Figure 1). For example, in areas where there is no presence of human beings, traps were installed randomly in vegetation. For areas with human presence, such as the technical offices’ of guard stations, traps were located inside and outside of buildings. For each sampling point, four different field collection techniques were used: HomeTrap-UIS, Torre Vigia-UIS, CDC, and Bg-Sentinel traps (Figure 2). HomeTrap-UIS and Torre Vigia-UIS are the types of sticky traps previously designed by our working group. These traps promote visual (red and black colors) and olfactory attraction (Sweetscent attractive in agar matrix) of insects and prevent insect escape by their particular design as small cardboard boxes (Figure 2) (Vidal et al. 2019). Bg-Sentinel trap is a folding cloth container that attracts mosquitoes by visual (black) and/or olfactory stimulus (Biogents 2017), whereas CDC uses blue light attraction (John W. Hock Company 2020); both traps suck mosquitoes inside with a fan (Biogents 2017, John W. Hock Company 2020). Two CDC and one Bg-Sentinel traps, both with Biogents Sweetscent attractive, were placed one night in each sampling point monthly, from 6:00 p.m. to 6:00 a.m. Six HomeTrap-UIS and six Torre Vigia-UIS traps were installed in each sampling point for one month. In summary, sampling effort by sampled point was 8,676 h/month, distributed in 24 h for CDC, 12 h for Bg-Sentinel, 4,320 h (24 h * 30 d * 6 traps) for HomeTrap-UIS trap, and 4,320 h for Torre Vigia-UIS trap.

**Figure 2.**
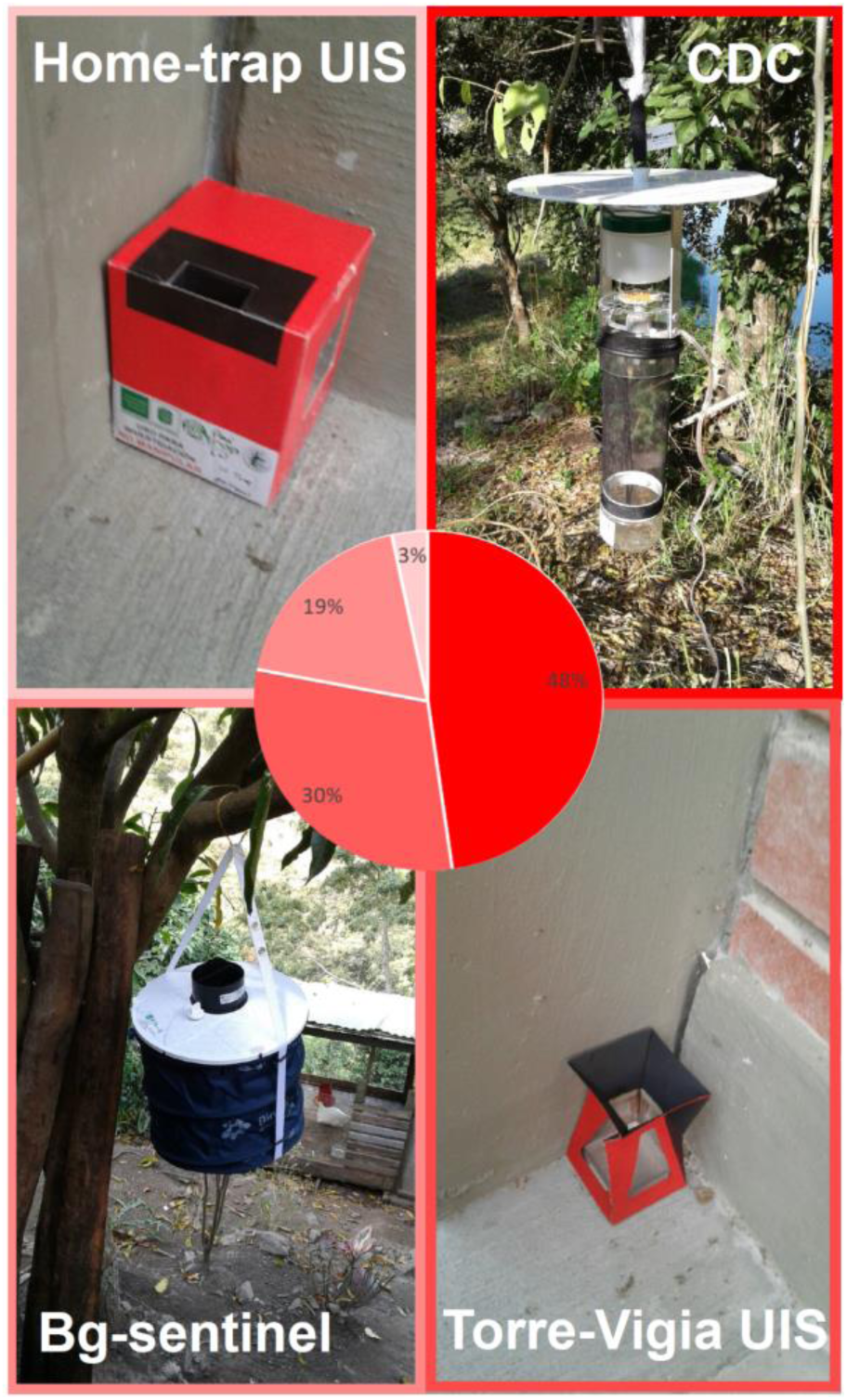
Relative sand fly amount collected by trap, around the artificial water reservoir Tona (Santander, Colombia) in 2017.

### 2.4 Specimen processing and identification

Collected sand flies were separated from other insects and placed on paper inside labeled Petri dishes for later processing. When well preserved in the traps, they were cleared by potassium hydroxide (10 – 20%) 12 – 24 h and neutralized with acetic acid (100%). Then, they were dyed with fuchsin acid for 15 min and washed with alcohol (70, 90, 100%, 10 min each). Finally, we fixed the specimens with Entellan or Canada balsam oils on glass slides. When specimens were poorly preserved, they were mounted on entomological card points. For species identification, structures of the head, thorax, and abdomen were used according to the taxonomic key proposed by Galati (2018).

### 2.7. Data analysis

The total number of sand flies collected was determined by type of sampling, species, month, and sampling site. Sand fly diversity in the Tona Reservoir was analyzed for the complete assemblage area. The amount of each species was determined according to the total number of individuals collected by species/morpho-species. The observed species richness (Sobs) corresponds to the cumulative number of species by month. To infer how much of the study area’s total expected sand fly richness was obtained in this study, we estimates cumulative curves of species with the Chao1 estimator and the amount-based coverage estimator (ACE), using the number of species collected by month as the sampling unit and EstimateS 9.1 program (Colwell 2000). To calculate the study area’s diversity, the Chao1 estimator focuses on the number of rare species (Magurran, 2004); Whereas, ACE estimator emphasizes the relevance of common species, i.e. species that are abundant or widely distributed spatially (Colwell and Coddington, 1994). Singletons and doubletons also provide information about rare species, considering the number of samples collected once and twice per month, respectively. Finally, to graph the turnover of the relative amount of species between months and places sampled, the statistical program Prisma 9.0 was used.

## 3. Results

268 sand flies were collected around the Tona Reservoir (Bucaramanga, Santander, Colombia) from February to November of 2017; 64.6% were females, and 35.4% were males (Table 1). Most specimens were collected with CDC (47.8%) and Torre Vigia-UIS (30.2%) traps (Table 2, Figure 2). 126 specimens (47%) could not be determined because the samples’ preservation status was poor; these samples came mostly from the sticky traps (30.2% Torre Vigia-UIS, 3.3% HomeTrap-UIS, 7.1% CDC, 6.3% Bg-sentinel) because specimens got stuck in the traps (Table 2). The remaining 53% of samples were identified to species/genus level. We found nine genera and 16 sand fly species, of which 12 were determined to species level (Figures 3 and 4). Two species correspond to new records for the Santander department: *Pi. youngi* (Feliciangeli and Murillo, 1987) and *Lu. ceferinoi* (Ortíz and Álvarez, 1963). *Pintomyia youngi*, *Pressatia camposi* (Rodríguez, 1950), and *Evandromyia dubitans* (Sherlock, 1962) presented the highest relative amount for four or more months (Figure 3b). In general, amount of these species was high: *Pi. youngi*: 17.9%; *Pr. camposi*: 10.4%, *Ev. dubitans*: 6.7%. Another relevant species collected around the reservoir was *Lu. gomezi.* This species presented a low relative amount (2.2%) and was collected for five months. The remaining species were relatively rare, exhibiting relative amounts from 0.7 to 2.6% and were collected in one or two months (Figure 3b).

**Figure 3.**
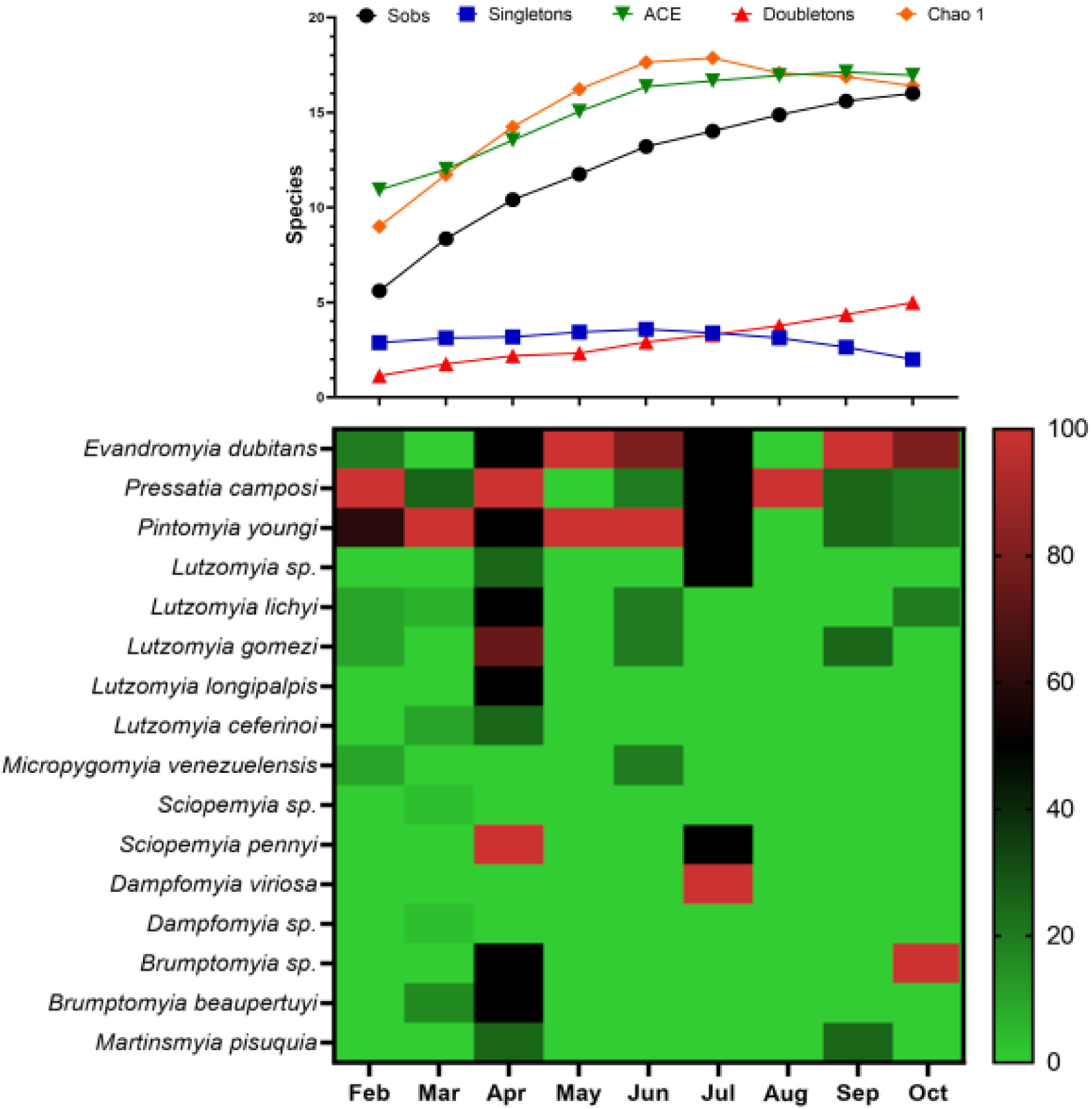
(a) Diversity estimator’s cumulative curves and (b) relative amount of sand fly species by month for the artificial water reservoir Tona (Santander, Colombia) in 2017.

**Figure 4.**
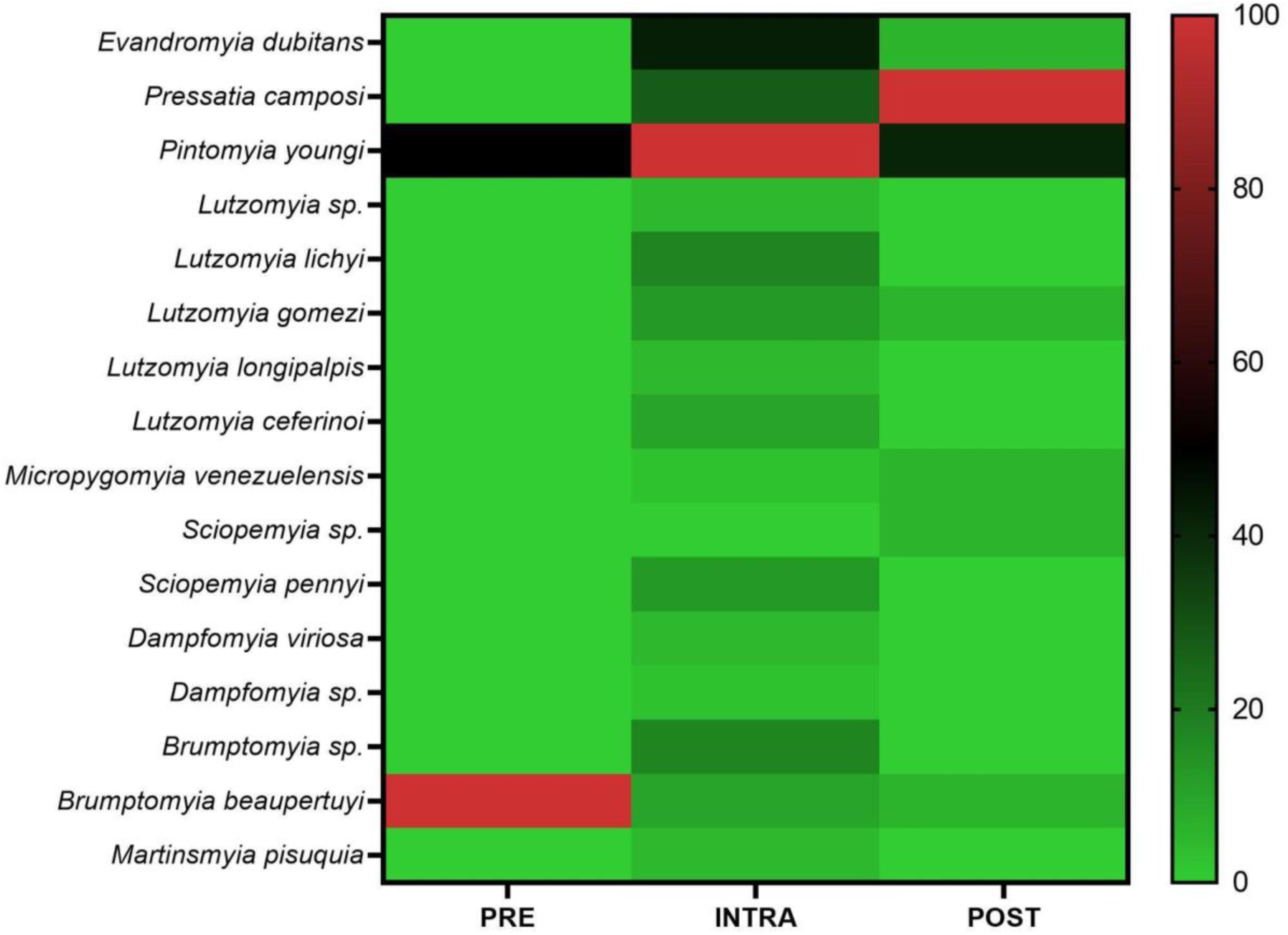
Spatial variation in relative sand fly amount across pre-, intra-, and post-Tona water reservoir areas (Santander, Colombia).

**Table 1.** Sand flies collected in the artificial water reservoir Tona (Santander, Colombia) in 2017.

| Species | Female | Male | Total |
| --- | --- | --- | --- |
| *Evandromyia dubitans* | 2 | 16 | 18 |
| *Pressatia camposi* | 25 | 3 | 28 |
| *Pintomyia youngi* | 42 | 6 | 48 |
| *Lutzomyia* sp. | 2 | 0 | 2 |
| *Lutzomyia lichyi* | 7 | 0 | 7 |
| *Lutzomyia gomezi* | 4 | 2 | 6 |
| *Lutzomyia longipalpis* | 0 | 2 | 2 |
| *Lutzomyia ceferinoi* | 3 | 1 | 4 |
| *Micropygomyia venezuelensis* | 2 | 0 | 2 |
| *Sciopemyia* sp. | 1 | 0 | 1 |
| *Sciopemyia pennyi* | 2 | 3 | 5 |
| *Dampfomyia viriosa* | 2 | 0 | 2 |
| *Dampfomyia* sp. | 1 | 0 | 1 |
| *Brumptomyia* sp. | 3 | 4 | 7 |
| *Brumptomyia beaupertuyi* | 1 | 6 | 7 |
| *Martinsmyia pisuquia* | 0 | 2 | 2 |
| Phlebotominae spp. | 76 | 50 | 126 |
| TOTAL | 173 | 95 | 268 |
| % | 64.6 | 35.4 | 100 |

**Table 2.**
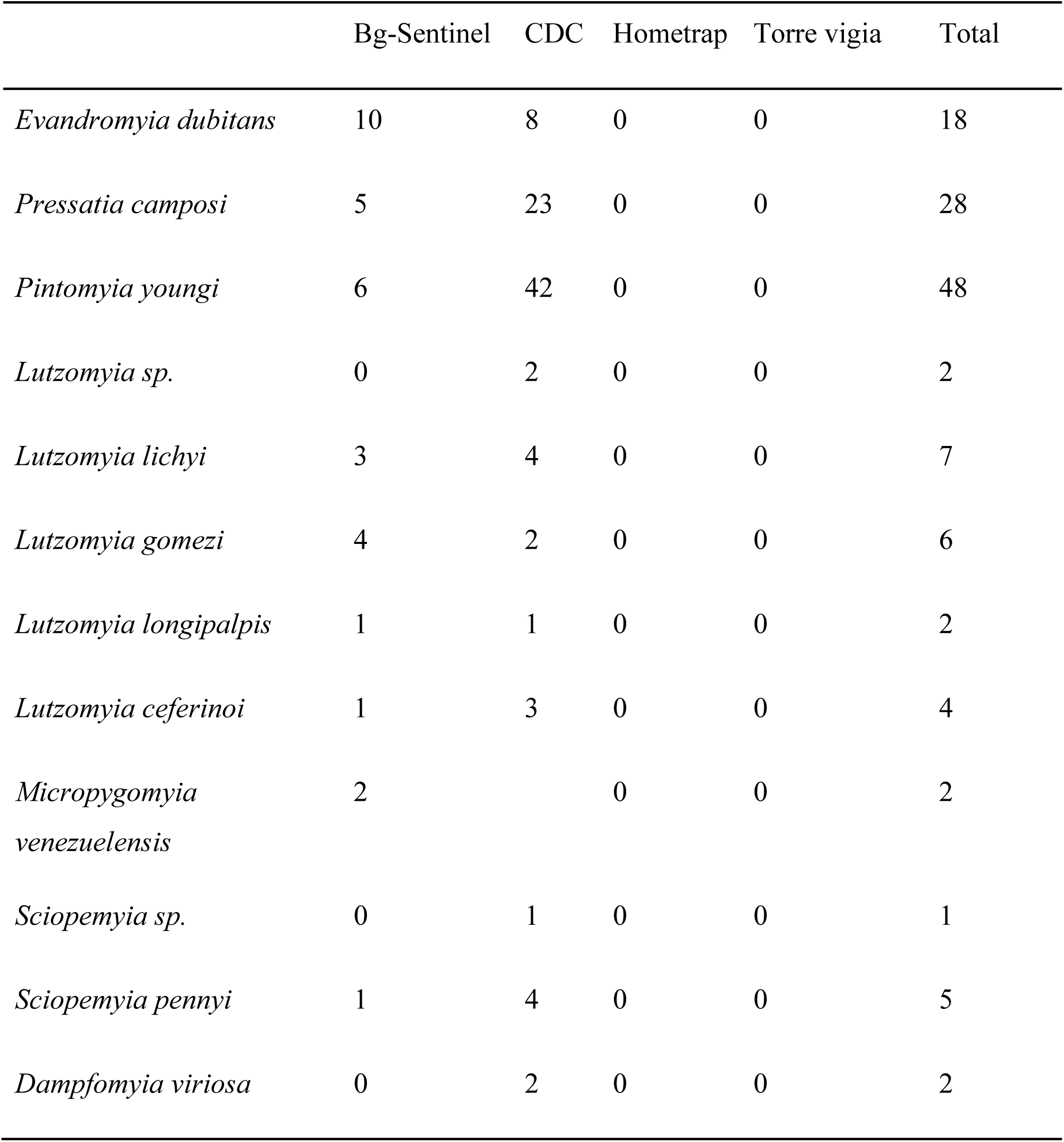
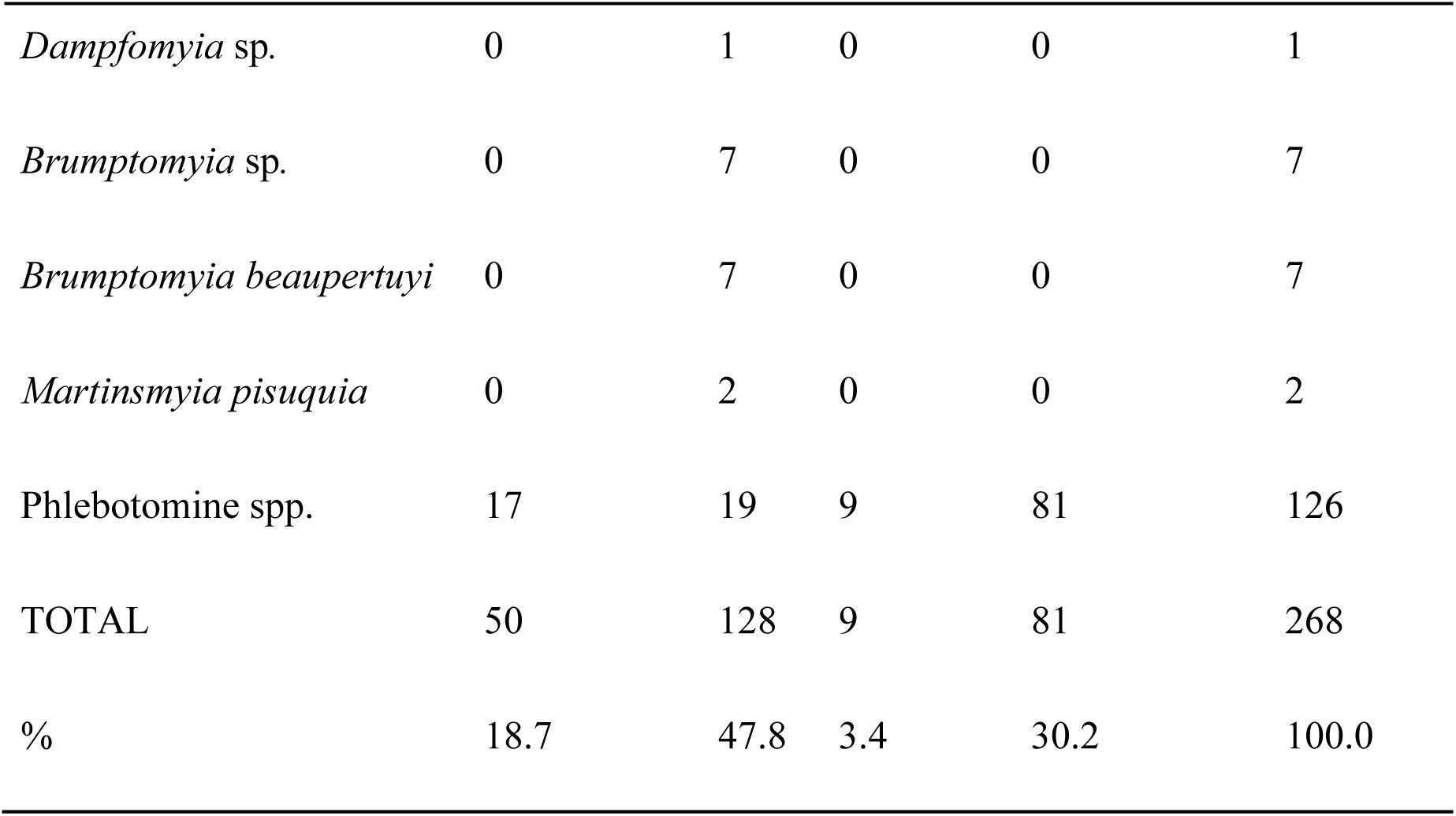
Sand fly species collected by trap in the artificial water reservoir Tona (Santander, Colombia) in 2017.

|  | Bg-Sentinel | CDC | Hometrap | Torre vigia | Total |
| --- | --- | --- | --- | --- | --- |
| *Evandromyia dubitans* | 10 | 8 | 0 | 0 | 18 |
| *Pressatia camposi* | 5 | 23 | 0 | 0 | 28 |
| *Pintomyia youngi* | 6 | 42 | 0 | 0 | 48 |
| *Lutzomyia sp.* | 0 | 2 | 0 | 0 | 2 |
| *Lutzomyia lichyi* | 3 | 4 | 0 | 0 | 7 |
| *Lutzomyia gomezi* | 4 | 2 | 0 | 0 | 6 |
| *Lutzomyia longipalpis* | 1 | 1 | 0 | 0 | 2 |
| *Lutzomyia ceferinoi* | 1 | 3 | 0 | 0 | 4 |
| *Micropygomyia venezuelensis* | 2 |  | 0 | 0 | 2 |
| *Sciopemyia sp.* | 0 | 1 | 0 | 0 | 1 |
| *Sciopemyia pennyi* | 1 | 4 | 0 | 0 | 5 |
| *Dampfomyia viriosa* | 0 | 2 | 0 | 0 | 2 |
| *Dampfomyia* sp. | 0 | 1 | 0 | 0 | 1 |
| *Brumptomyia* sp. | 0 | 7 | 0 | 0 | 7 |
| *Brumptomyia beaupertuyi* | 0 | 7 | 0 | 0 | 7 |
| *Martinsmyia pisuquia* | 0 | 2 | 0 | 0 | 2 |
| Phlebotomine spp. | 17 | 19 | 9 | 81 | 126 |
| TOTAL | 50 | 128 | 9 | 81 | 268 |
| % | 18.7 | 47.8 | 3.4 | 30.2 | 100.0 |

Species accumulation curves allowed us to infer that the sampling made for the Tona Reservoir in 2017 was representative of the area’s corresponding diversity. Curves of ACE and Chao 1 estimator and the observed species (Sobs) curve tended to reach the asymptote. This indicated that a minimum sampling effort is required to know the complete sand fly fauna of the Tona Reservoir. The singletons (rare species) curve also shows that an asymptote has already been reached, while the curve of doubletons shows that it needs more sampling effort to reach an asymptote (Figure 3a). The highest richness of species was collected in April (12 spp). This month, the relative amount of sand flies was also the highest of the sampled period (February to November 2017).

In consonance with the sampling effort, the intra-Tona Reservoir exhibited an outstanding diversity, presenting the highest richness of species (15 spp.) and the largest number of samples (84.3%, Figure 4). Sampling effort by area was highly variable due to the differential number of sand fly sampling points (Figure 1). Sampling effort by area was: 86,760 h/month for the intra-Tona Reservoir (ten sampling points), 26,028 h/month for the post-Tona Reservoir (three), and 8,676 h/month for the pre-Tona Reservoir (one). *Pi. youngi*, the most abundant species in this study, was found in pre-, intra-, and post-Tona Reservoir areas. *Pressatia camposi* and *Ev. dubitans* were recorded around post– and intra-Tona Reservoir areas, respectively. It is worth mentioning that only *Pi. youngi* and *Brumptomyia beaupertuyi* were recorded around the pre-Tona Reservoir area (Figure 4).

## 4. Discussion

Here, we characterized sand fly biodiversity around the Tona Reservoir in 2017 and compared performance in the field of two sticky traps previously designed by our working group (Torre Vigia-UIS, HomeTrap-UIS) with commonly used traps of common use (CDC, Bg-Sentinel traps). The Tona Reservoir was built in 2016 in a tropical wet forest –bh-PM (Rincón et al. 2019)– of Santander (Colombia). Our study showed that sand fly biodiversity in this water reservoir was high, with numbers that represent 8% and 30% of the sand fly fauna of Colombia and Santander, respectively. Among species of medical relevance collected in the area, *Pi. youngi* seems particularly relevant because of its wide spatiotemporal distribution. Torre Vigia-UIS emerged as a valuable tool for vector control because it was very efficient for collecting samples in the Tona water reservoir.

The natural features of the study area provided conditions for the high diversity of the Tona Reservoir. Another similar study was performed in Sogamoso dam, which is located in a tropical wet forest of Santander; this study reported 21 spp. in 2016-2017 (Gómez-Vargas and Zapata-Úsuga 2022), comparable to our findings. The highest diversity was collected around the most intervened area, the Intra-Tona reservoir, and in the rainy season. Sand fly diversity was three times more diverse around the Intra-Tona (15 spp.) than around the Post-Tona reservoir, but this difference is probably related to a differential sampling effort. Instead of spatial differences, the rainy season’s climatic conditions were relevant for the Tona Reservoir’s sand fly diversity. Bh-PM is a tropical wet forest with bimodal rainfalls (Rincón et al. 2019). Most species were collected in April, one of the months of the rainy season, and these species were collected in the largest relative amount of the sampled period (Figure 3b). In general, our curves of biodiversity estimations (Figure 3a) lead us to infer that we were close to recording the complete sand fly fauna of the Tona Reservoir. It is worth mentioning that our diversity estimations were not affected by a miscount of morphospecies. Morphoespecies could affect the counts of several species, for instance, by confusing different specimens of the same species as two separate species. Morphospecies studied here were counted as just one species because we could make full morphological comparisons among these specimens, as well as between them and their closest species.

Considering our new records, Santander has 53 sand fly species. The last review of sand fly fauna for Colombia reported 49 species in Santander (Bejarano and Estrada 2016). Recent research in Sogamoso dam added *Lu. sanguinaria* y *Lu. strictivilla* to these records (Gómez-Vargas and Zapata-Úsuga 2022). Here, we added *Lu. ceferinoi* and *Pi. youngi* to sand fly records for Santander. *Lutzomyia ceferinoi* has records for Colombia and Venezuela (Bejarano and Estrada 2016). The *Pi. youngi* record was an unexpected finding for Santander because this department is far from its known distribution area. Its known records came from Antioquia, Caldas, and Valle del Cauca departments in the northern Andean western mountains (Bejarano and Estrada 2016). This species has also been recorded in Venezuela and Costa Rica (Feliciangeli and Murillo 1987). *Pintomyia youngi* is a vector of *L. braziliensis* mostly associated with coffee farms (Contreras-Gutiérrez et al. 2014) and, in the Tona Reservoir, might provide a continuous leishmaniasis risk because of its broad spatiotemporal distribution. Other species of medical relevance found in the Tona Reservoir were *Lu. longipalpis* and *Lu. gomezi*.

Among the four sampling methods evaluated, CDC and Torre Vigia-UIS both excelled. This is the first time the Torre Vigia-UIS performance has been evaluated in the field. CDC was undoubtedly the best sampling method for our study because this method reached a large sample collection in a few hours (24 h by sampling point). This finding is congruent with previous evidence about CDC vs. box trap performance (Kweka et al. 2013). The Torre Vigia-UIS trap was left for a long time in each sample point (4320 h), and most of the specimens collected with this sticky trap had poor preservation status, disabling any taxonomic determination. Therefore, we do not recommend this trap for biodiversity studies because this kind of study requires fast characterization of faunas and well-preserved material. However, Torre Vigia-UIS has other features that make it a valuable tool for vector control. This trap is very cheap, with a non-commercial value of $160 US for a pack of 60 traps –this price is similar to just one CDC trap–. The Torre Vigia-UIS trap is easy and fast to assemble, it is safe and eco-friendly to use, it does not use batteries or any recharging procedure, and it does not require any maintenance throughout its operation (Vidal et al. 2019). Considering that most communities affected by leishmaniasis exhibit multidimensional poverty, providing a simple but efficient vector trap might provide more options for integrated disease management.

## Acknowledgments

We thanks to Ellen Cannon for assisting us in improving the English language of our manuscript. L.R-C. thanks, Vicerrectoría de Investigaciones of the University Industrial of Santander, for support through the Postdoctoral fellow program.

